# Schizophrenia risk and reproductive success: A Mendelian randomization study

**DOI:** 10.1101/357673

**Authors:** Rebecca B Lawn, Hannah M Sallis, Amy E Taylor, Robyn E Wootton, George Davey Smith, Neil M Davies, Gibran Hemani, Abigail Fraser, Ian S Penton-Voak, Marcus R Munafò

## Abstract

Schizophrenia is a debilitating and heritable mental disorder associated with lower reproductive success. However, the prevalence of schizophrenia is stable over populations and time, resulting in an evolutionary puzzle: how is schizophrenia maintained in the population given its apparent fitness costs? One possibility is that increased genetic liability for schizophrenia, in the absence of the disorder itself, may confer some reproductive advantage. We assessed the correlation and causal effect of genetic liability for schizophrenia with number of children and age at first birth using data from the Psychiatric Genomics Consortium and UK Biobank. Linkage disequilibrium score regression showed little evidence of genetic correlation between genetic liability for schizophrenia and number of children (rg=0.002, *p*=0.84) or age at first birth (rg=-0.007, *p*=0.45). Mendelian randomization indicated no robust evidence of a causal effect of genetic liability for schizophrenia on number of children (mean difference: 0.003 increase in number of children per doubling in the natural log odds ratio of schizophrenia risk, 95% CI: −0.003 to 0.009, *p*=0.39) or age at first birth (−0.004 years lower age at first birth, 95% CI: −0.043 to 0.034, *p*=0.82). These results suggest that increased genetic liability for schizophrenia does not confer a reproductive advantage.

## Introduction

Schizophrenia is a severe, debilitating mental disorder that is substantially heritable (1). The prevalence of schizophrenia remains stable over populations and time, and yet is associated with lower reproductive success for those diagnosed (1–5). This creates an evolutionary puzzle: how is schizophrenia maintained in the population despite apparent negative selection? Multiple theories have been proposed to explain this paradox (3,6,7). One is mutation-selection balance, which suggests that selection against detrimental variants is counteracted by the continuous occurrence of new mutations (8,9). Another is that effects over many common genetic variants are individually too weak to be under negative selection (1,8,10).

Another popular theory is that stabilizing selection operates, where the optimum fitness level for a trait is approximately at the mean of the trait and fitness declines along a normal distribution on either side of this optimum (3,6,11,12). A related hypothesis is that schizophrenia-related traits may demonstrate ‘cliff-edge’ effects on fitness, so that fitness increases with increased expression of the trait until a threshold, where increased expression then results in a steep decline in fitness for some individuals (1,12). Some have suggested that this peak occurs at levels of symptoms that could result in a diagnosis of schizophrenia, with a reproductive advantage among healthy individuals with an increased genetic liability for the disorder (in the absence of the disorder itself) compensating for the lower reproductive success of those with the disorder itself (1,4,12–14). Behaviourally, it is possible that higher genetic liability for schizophrenia may be associated with attractive traits (e.g., creativity) and therefore also with greater number of children (4,13). For example, schizotypy, a personality measure of schizophrenia-proneness, has been shown to be associated with creativity, short term mating interest and mating success (4,13,15), while genetic liability for schizophrenia is associated with increased risk of unprotected sex (16).

Relatives of people with schizophrenia are assumed to have an intermediate level of genetic liability for the highly heritable disorder (17). Studies into whether cliff-edge fitness maintains the prevalence of schizophrenia have therefore largely focused on family studies. However, despite extensive research, there is no clear evidence of increased fecundity in relatives of individuals with schizophrenia (2,7,17). Del Giudice argued that family studies underestimate the reproductive benefits of schizophrenia-proneness in the general population (17). He highlights that relatives not only share genetic liability for schizophrenia but also their environments, which may hinder fitness and result in apparent negative selection (17). It is therefore important to investigate a potential reproductive advantage of schizophrenia-proneness in the wider population, rather than relying on family studies alone. Moreover, it is important to investigate causal relationships between schizophrenia risk and reproductive success, rather than relying on observational methods previously used, which do not support strong causal inference due to bias from residual confounding and reverse causation (18). These family studies also suggest that optimum fitness could occur before the appearance of symptoms that might result in a diagnosis of schizophrenia.

Recent developments in genetic epidemiology mean that it is now possible to investigate the effects of genetic liability for schizophrenia in the wider population. A genome-wide association study (GWAS) identified 128 independent genetic variants from 108 loci associated with schizophrenia that explained approximately 3.4% of the observed variation in schizophrenia risk (19). These variants have been used to show that genetic risk for schizophrenia (using a risk score comprising these individual variants) is positively associated with creativity (20). In the context of reproductive success, earlier age at first birth, likely resulting in a longer reproductive period, can be used as an indicator of this (8,21,22). However, evidence for associations between genetic liability for schizophrenia and age at first birth is mixed. Higher genetic liability for schizophrenia was found for those with a young (e.g., below 20 years) age at first birth compared to those with an intermediate age at first birth (23,24). Another study found no clear evidence for linear or quadratic associations between a genetic liability for schizophrenia and age at first birth (8). Two previous studies also used schizophrenia-associated variants to investigate whether genetic liability for schizophrenia is associated with number of children, but results were again inconclusive, perhaps due to limited power (8,25). The studies showed estimates in the direction of a reproductive advantage, but confidence intervals are typically wide and consistent with no effect (8,25). Nevertheless, these studies demonstrate how genetic liability for schizophrenia can be measured in the wider population.

We applied a range of methods with roots in genetic epidemiology to test part of the cliff-edge hypothesis. For our main analysis, we examine whether increasing genetic liability for schizophrenia increases reproductive success in a largely post-reproductive population-based sample which is not selected on schizophrenia status, and therefore includes very few cases. This linear increase is predicted for part of cliff-edge fitness where a reproductive advantage among healthy individuals with higher genetic liability for the disorder compensates for lower reproductive success of those with the disorder itself. Additionally, given suggestions from family studies that there may be a fitness decline of healthy individuals with high genetic liability for the disorder, we conducted sensitivity analyses to investigate a possible non-linear relationship where at very high levels of the score there is lower fecundity (2,7,17).

For our main analysis, we calculated genetic correlations using LD-score regression between genetic liability for schizophrenia and reproductive success, measured as number of children and age at first birth. Furthermore, we used genetic variants associated with schizophrenia within a Mendelian Randomization (MR) framework to estimate the causal effect of genetic liability for schizophrenia and these measures of reproductive success. MR uses single nucleotide polymorphisms (SNPs), which are assigned at conception and are mostly independent from other variants or environments. MR therefore overcomes some limitations of observational studies previously used to investigate this evolutionary paradox, by reducing bias from confounding and reverse causation (18). Finally, we estimated the effect of genetically-predicted educational attainment on number of children and age at first birth as a positive control, where the direction of results is known, with the same outcome datasets used for our schizophrenia analysis, given that higher genetically-predicted education is known to be associated with fewer children (25–29). Our results show little evidence that schizophrenia and reproductive success are genetically correlated in the general population, or that high liability for schizophrenia affects fecundity, suggesting that the sustained prevalence of schizophrenia in the population is not due to cliff-edge fitness.

## Methods

### Exposure data

We used independent single nucleotide polymorphisms (SNPs) associated with schizophrenia (*p* < 5 × 10^−8^) from the Psychiatric Genomics Consortium GWAS (*N* = 36,989 cases and 113,075 controls) (19). The 128 SNPs identified explained approximately 3.4% of the observed variance in schizophrenia. A total of 101 SNPs remained due to availability in UK Biobank, availability of proxies, and meeting exclusion criteria (see Supplementary Text). Odds ratios (ORs) and standard errors (SE) for the 101 SNP and schizophrenia associations were recorded using GWAS data for Europeans only (30). The final 101 SNPs and effect estimates for schizophrenia genetic variants are listed in Supplementary Table 1.

For educational attainment, we used SNPs associated with educational attainment (*p* < 5 × 10^−8^) from a recent GWAS by the Social Science Genetic Association Consortium (31). As the GWAS conducted a replication in UK Biobank, effect estimates from the pooled sex analysis of the discovery sample were used to avoid sample overlap. Sixty-seven SNPs were available in UK Biobank data and met exclusion criteria. Effect estimates used for educational attainment genetic variants are listed in Supplementary Table 2.

### Outcome data

The exposure associated SNPs described above were extracted from UK Biobank to derive SNP-outcome associations for our outcome data. Extraction was done using PLINK (v2.00) and best guess algorithms for determining alleles (full genotyping information below).

#### Sample

UK Biobank is a population-based health research resource consisting of approximately 500,000 people, aged between 38 years and 73 years, who were recruited between the years 2006 and 2010 from across the UK (32). Particularly focused on identifying determinants of human diseases in middle-aged and older individuals, participants provided a range of information (such as demographics, health status, lifestyle measures, cognitive testing, personality self-report, and physical and mental health measures) via questionnaires and interviews; anthropometric measures, BP readings and samples of blood, urine and saliva were also taken. A full description of the study design, participants and quality control (QC) methods has been published (33).

#### Genotyping information in UK Biobank

The full data release contains the cohort of successfully genotyped samples (*N* = 488,377). A total of 49,979 individuals were genotyped using the UK BiLEVE array and 438,398 using the UK Biobank axiom array. Pre-imputation QC, phasing and imputation are described elsewhere (34). In brief, prior to phasing, multiallelic SNPs or those with minor allele frequency (MAF) ≤1% were removed. Phasing of genotype data was performed using a modified version of the SHAPEIT2 algorithm (35). Genotype imputation to a reference set combining the UK10 K haplotype and HRC reference panels (36) was performed using IMPUTE2 algorithms (37). The analyses presented here were restricted to autosomal variants within the HRC site list using a graded filtering with varying imputation quality for different allele frequency ranges. Therefore, rarer genetic variants are required to have a higher imputation INFO score (Info>0.3 for MAF >3%; Info>0.6 for MAF 1–3%; Info>0.8 for MAF 0.5–1%; Info>0.9 for MAF 0.1–0.5%) with MAF and Info scores having been recalculated on an in house derived ‘European’ subset. Individuals with sex-mismatch (derived by comparing genetic sex and reported sex) or individuals with sex-chromosome aneuploidy were excluded from the analysis (*N* = 814). We restricted the sample to individuals of white British ancestry who self-report as “White British” and who have very similar ancestral backgrounds according to the PCA (*N* = 409,703), as described by Bycroft (34). Estimated kinship coefficients using the KING toolset (38) identified 107,162 pairs of individuals (34). An in-house algorithm was then applied to this list and preferentially removed the individuals related to the greatest number of other individuals until no related pairs remain. These individuals were excluded (*N* = 79,448). Additionally, 2 individuals were removed due to them relating to a very large number (>200) of individuals. Quality Control protocol is described elsewhere (39).

#### Outcome measures

We derived number of children and age at first birth similarly to previous analyses in UK Biobank (28). Participants were either asked how many children they had given birth to or how many children they had fathered. We further derived a binary variable to indicate if participants were childless or not (childlessness coded as 1). Age at first birth was only measured in females in UK Biobank, with participants asked: “How old were you when you had your first child?”. Although no age restrictions were applied in analyses, the nature of UK Biobank data meant that participants were aged towards the end of their reproductive lives.

### Data analysis

We used LD score regression (40,41) to calculate the genome-wide genetic correlation (r_g_) between schizophrenia liability or predicted educational attainment and number of children and age at first birth. Number of children and age at first birth genome-wide associations were conducted using linear regression, implemented in PLINK v2.00 through the MRC IEU GWAS pipeline (42). In this, we adjusted for the top 10 principal components. For number of children analysis, age and sex were also included as covariates. We then filtered results on MAF (>0.01) and imputation quality (>0.8) separately.

In MR analyses, data were harmonized to ensure that the effect of the SNP on the exposure and the SNP on the outcome corresponded to the same allele. The increasing allele for schizophrenia liability and educational attainment was used. Associations for exposure SNPs and number of children and age at first birth were then calculated in R, fitting the same covariates as listed above. Effect sizes for each analysis are listed in Supplementary Tables 1 and 2. SNP-exposure and SNP-outcome data were combined using an inverse variance weighted (IVW) approach which is analogous to a weighted regression of SNP-outcome coefficients on SNP-exposure coefficients with the intercept constrained to zero (43), and further assumes all variants are valid instruments or allows pleiotropy to be balanced across instruments when using the random effect (44) with Cochran’s Q providing a measure of any overdispersion (see Supplementary Text).

The IVW effect estimate will only be consistent if all genetic variants in the analysis are valid. Weighted median, mode-based estimator and MR-Egger regression are complementary approaches that can be used to investigate the impact of invalid instruments on our effect estimates. The weighted median estimates a consistent effect estimate if at least 50% of the instruments are valid (45). The mode-based estimator provides a consistent effect estimate when the largest number of similar individual-instrument estimates come from valid instruments, even if the majority are invalid (46). A tuning parameter of 0.5 was set for mode-based estimator analysis. One of the main assumptions underpinning MR is that of no horizontal pleiotropy (i.e., no direct effect of the genetic variant on the outcome that does not act through the exposure) (47). MR-Egger regression analysis can be used to further investigate this; MR-Egger does not constrain the intercept to zero and the intercept term therefore estimates overall directional pleiotropy (48). We calculated F statistics (mean of the squared SNP-exposure association divided by the squared standard error for SNP-outcome association) to indicate the strength of instrument, and I^2^GX statistics to assess the suitability of MR-Egger (above 0.9 is desired) (47). Analysis was repeated after removing the few schizophrenia cases in our sample. All analysis was also conducted with SNP-outcome associations additionally adjusted for genotype array.

MR results were multiplied by 0.693 to represent the causal estimate per doubling in odds of schizophrenia risk (49). For childlessness as an outcome, all MR results were multiplied by 0.693 on the log-odds scale, and then exponentiated. The reported estimates therefore indicate the effect of doubling the odds of schizophrenia on the odds of childlessness. The effect of education on childlessness were converted to ORs by exponentiating log ORs.

As an illustration of shape of the schizophrenia liability-fecundity relationship, we created an additive unweighted genetic score for schizophrenia liability in UK Biobank. The score was created in R (version 3.2.0), with missing SNP data replaced with the mean value for that SNP across individuals. We then divided this score into quintiles and plotted this against mean number of children and mean age at first birth. As sensitivity analysis to assess if there was any decline in fitness within our sample at very high levels of genetic liability, we conducted a series of linear regressions in Stata (version: MP 15.1) between the genetic score and number of children, systematically removing cumulative centiles from the maximum. This analysis included adjustment for the top 10 principal components and was repeated after removing the few schizophrenia cases in our sample. Similarly, to further investigate a possible peak in fitness at high genetic liability for schizophrenia, we conducted quadratic regression analysis of the schizophrenia genetic score and number of children (adjusted for the top 10 principal components and also additionally adjusted for sex and age at assessment).

Analysis scripts are available on GitHub (50).

## Results

In our sample, from UK Biobank, there were more females than males, a majority had children, and a minority had college or university degree qualifications (Supplementary Table 3). The mean age was 56.9 years (SD: 8.0), the mean number of children was 1.8 (SD: 1.2), and the mean age at first birth was 25.4 years (SD: 4.5). The mean years of education was 13.3 (SD: 4.4).

### Genetically predicted educational attainment

We found a modest negative genetic correlation between educational attainment associated variants and number of children (*r*_g_ = −0.35, *p* = 8.57 × 10−^−41^) and a strong positive genetic correlation between educational attainment associated variants and age at first birth (*r*_g_ = 0.81, *p* < 5×10^−41^) (Table 1).

**Table 1.** Genetic correlations of genetic liability for schizophrenia and genetically predicted educational attainment on number of children and age at first birth using LD score regression.

|  | No. of children<sup>a</sup> |  |  | Age at first birth<sup>b</sup> |  |  |
| --- | --- | --- | --- | --- | --- | --- |
|  | *r<sub>g</sub>* | *se* | *P* | *r<sub>g</sub>* | *se* | *P* |
| Genetic liability for schizophrenia<sup>c</sup> | 0.002 | 0.008 | 0.837 | -0.007 | 0.009 | 0.445 |
| Genetically predicted educational attainment<sup>d</sup> | -0.347 | 0.026 | $8.57 \times 10^{-41}$ | 0.805 | 0.019 | $<5 \times 10^{-41}$ |

<sup>a</sup> Number of children data from UK Biobank ( $N = 333,628$ ); <sup>b</sup> Age at first birth data from UK Biobank ( $N = 123,310$ ); <sup>c</sup> Schizophrenia data from the Psychiatric Genomics Consortium GWAS ( $N = 36,989$ cases and $113,075$ controls) (Consortium, 2014); <sup>d</sup> Educational attainment from the Social Science Genetic Association Consortium GWAS ( $N = 283,723$ ).

Educational attainment variants showed a mean F statistic (strength of instrument) of 33.23, with above 10 indicating acceptable levels of relative bias (<10%) (44,51). We applied muliple MR methods with IVW results reported throughout the text, and other methods only when not consistent. We found that educational attainment had a negative effect on number of children (mean difference: −0.16, 95% confidence interval [CI]: −0.21 to −0.12, *p* = 3.63 × 10^−10^ per year increase in educational attainment) and a positive effect on age at first birth (mean difference 2.68, 95% CI: 2.40 to 2.95, *p* <5×10^−14^) per year increase in educational attainment) (Table 2). We also found an effect of increased education on increased likelihood of being childless (odds ratio [OR]: 1.38, 95% CI: 1.29 to 2.00, *p* = 1.60 × 10–^−14^ per year increase in educational attainment). Results for all educational attainment analysis with genotype array included as a covariate in our outcome summary statistics are presented in Supplementary Tables 4 and 5.

**Table 2.** Estimates of the causal effect of genetic liability for schizophrenia and genetically predicted educational attainment on number of children, age at first birth and childlessness using inverse variance weighted, mode-based estimator, MR-Egger and weighted median Mendelian randomization approaches.

|  | No. of children<sup>b</sup> | Age at first birth<sup>c</sup> | Childlessness<sup>d</sup> |
| --- | --- | --- | --- |
| Method | $\beta$ (95% CI), *P* | | OR (95% CI), *P* |
| **Genetic liability for schizophrenia: 101 SNPs<sup>a</sup>** |  |  |  |
| Inverse Variance Weighted | 0.003 (-0.003, 0.009), 0.39 | -0.004 (-0.043, 0.034), 0.82 | 0.998 (0.985, 1.012), 0.79 |
| MR-Egger intercept | -0.001 (-0.004, 0.001), 0.29 | -0.016 (-0.031, -0.001), 0.04 | 0.998 (0.993, 1.004), 0.55 |
| Mr-egger slope | 0.020 (-0.013, 0.053), 0.23 | 0.214 (0.007, 0.420), 0.04 | 1.019 (0.950, 1.094), 0.59 |
| Weighted median | 0.006 (-0.004, 0.015), 0.23 | 0.023 (-0.042, 0.089), 0.49 | 0.995 (0.975, 1.016), 0.65 |
| Simple mode-based estimator | 0.020 (-0.014, 0.055), 0.25 | 0.067 (-0.196, 0.331), 0.62 | 0.988 (0.912, 1.070), 0.76 |
| Weighted mode-based estimator | 0.020 (-0.012, 0.052), 0.22 | 0.060 (-0.175, 0.294), 0.62 | 0.992 (0.924, 1.065), 0.83 |
| **Genetically predicted educational attainment: 67 SNPs<sup>e</sup>** |  |  |  |
| Inverse Variance Weighted | -0.162 (-0.206, -0.118), $3.63 \times 10^{-10}$ | 2.677 (2.401, 2.952), $<5 \times 10^{-14}$ | 1.589 (1.446, 1.746), $1.60 \times 10^{-14}$ |
| MR-Egger intercept | 0.004 (0.001, 0.008), $2.50 \times 10^{-02}$ | -0.031 (-0.054, -0.008), $9.52 \times 10^{-03}$ | 0.990 (0.982, 0.997), 0.010 |
| Mr-egger slope | -0.391 (-0.595, -0.187), $2.99 \times 10^{-04}$ | 4.348 (3.069, 5.628), $4.12 \times 10^{-09}$ | 2.812 (1.813, 4.362), $1.38 \times 10^{-05}$ |
| Weighted median | -0.206 (-0.276, -0.135), $2.93 \times 10^{-07}$ | 2.828 (2.387, 3.270), $<5 \times 10^{-14}$ | 1.567 (1.343, 1.829), $3.00 \times 10^{-07}$ |
| Simple mode-based estimator | -0.253 (-0.5107, 0.005), 0.06 | 3.454 (1.938, 4.969), $3.18 \times 10^{-05}$ | 1.474 (0.884, 2.457), 0.14 |
| Weighted mode-based estimator | -0.249 (-0.478, -0.020), 0.04 | 1.649 (0.303, 2.995), $1.92 \times 10^{-02}$ | 1.513 (0.952, 2.404), 0.085 |

<sup>a</sup> Schizophrenia genetic data from the Psychiatric Genomics Consortium GWAS (N= 36,989 cases and 113,075 controls); <sup>b</sup> Number of children data from UK Biobank (N = 318,921 – 335,758 for genetic liability of schizophrenia analysis and 268,658 – 335,758 for educational attainment analysis). Schizophrenia results were multiplied by 0.693 to represent the estimate per doubling in odds of the exposure; <sup>c</sup> Age at first birth data from UK Biobank (N = 117,844 – 124,093 for genetic liability of schizophrenia analysis and 99,317 – 124,093 for education analysis). Schizophrenia results were multiplied by 0.693 to represent the estimate per doubling in odds of the exposure; <sup>d</sup> Childlessness data from UK Biobank (N = 318,921 – 335,758 for schizophrenia genetic liability of schizophrenia analysis and 268,658 – 335,758 for educational attainment analysis). Childlessness was coded as 1. Results were converted to ORs for schizophrenia by multiplying log ORs by 0.693 and then exponentiating to represent the OR per doubling in odds of the binary exposure. Results were converted to ORs for educational attainment by exponentiating log ORs; <sup>e</sup> Educational attainment from the Social Science Genetic Association Consortium GWAS (N = 283,723). It should be noted that the $I^2_{GX}$ statistic for an unweighted
MR-Egger regression was 0.33 for educational attainment and 0.20 for genetic liability of schizophrenia, which is deemed too low to conduct a SIMEX adjustment, and MR-Egger results should be treated with caution (47).

### Genetic liability for schizophrenia

Using LD score regression, we found little evidence of genetic correlations (*r_g_*) between schizophrenia liability and number of children (*r_g_*=0.002, p=0.84) and age at first birth (*r_g_*=- 0.007, p=0.45) (Table 1).

The mean F statistic for schizophrenia genetic liability was 35.15. There was little evidence that higher genetic liability for schizophrenia increased number of children (mean difference: 0.003 increase in number of children per doubling in the natural log OR of schizophrenia libility, 95% CI: −0.003 to 0.009, *p* = 0.39) or decreased age at first birth (−0.004 years lower age at first birth, 95% CI: −0.043 to 0.034, *p* = 0.82) (Table 2). We further tested childlessness as an outcome and found no strong evidence of an effect of genetic liability for schizophrenia on childlessness (Table 2). We repeated the MR analysis after removing the few schizophrenia cases in our sample (maximum *N* = 207) with no clear change in results. Results for these analyses with genotype array included as a covariate in our outcome summary statistics are presented in Supplementary Tables 4 and 5.

Our sensitivity analysis investigating a possible non-linear relationship is presented in Figures 1 and 2, showing the mean number of children and mean age at first birth for quintiles of an unweighted additive genetic score for schizophrenia liability. Although these figures are somewhat suggestive of a non-linear relationship between a genetic score for schizophrenia liability and mean age at first birth, there is little evidence of heterogeneity across values of the schizophrenia score. Further sensitivity analysis investigating whether there is lower fecundity at very high levels of the score were suggestive of a decline in fitness with high genetic liability. A series of regressions between the genetic score and number of children, systematically removing cumulative centiles from the maximum, suggested that estimates become slightly stronger with increased trimming of the score, although there is little statistical support for this pattern (Table 3). This analysis was repeated after removing the few schizophrenia cases in UK Biobank (maximum *N* = 207), which did not alter these results (Supplementary Table 6). Quadratic regression of the schizophrenia genetic score and number of children similarly showed a slight peak in fitness at intermediate levels of the genetic liability, particularly for females, but again with little statistical support (Supplementary Table 7 and Supplementary Figures 1–3).

**Figure 1.**
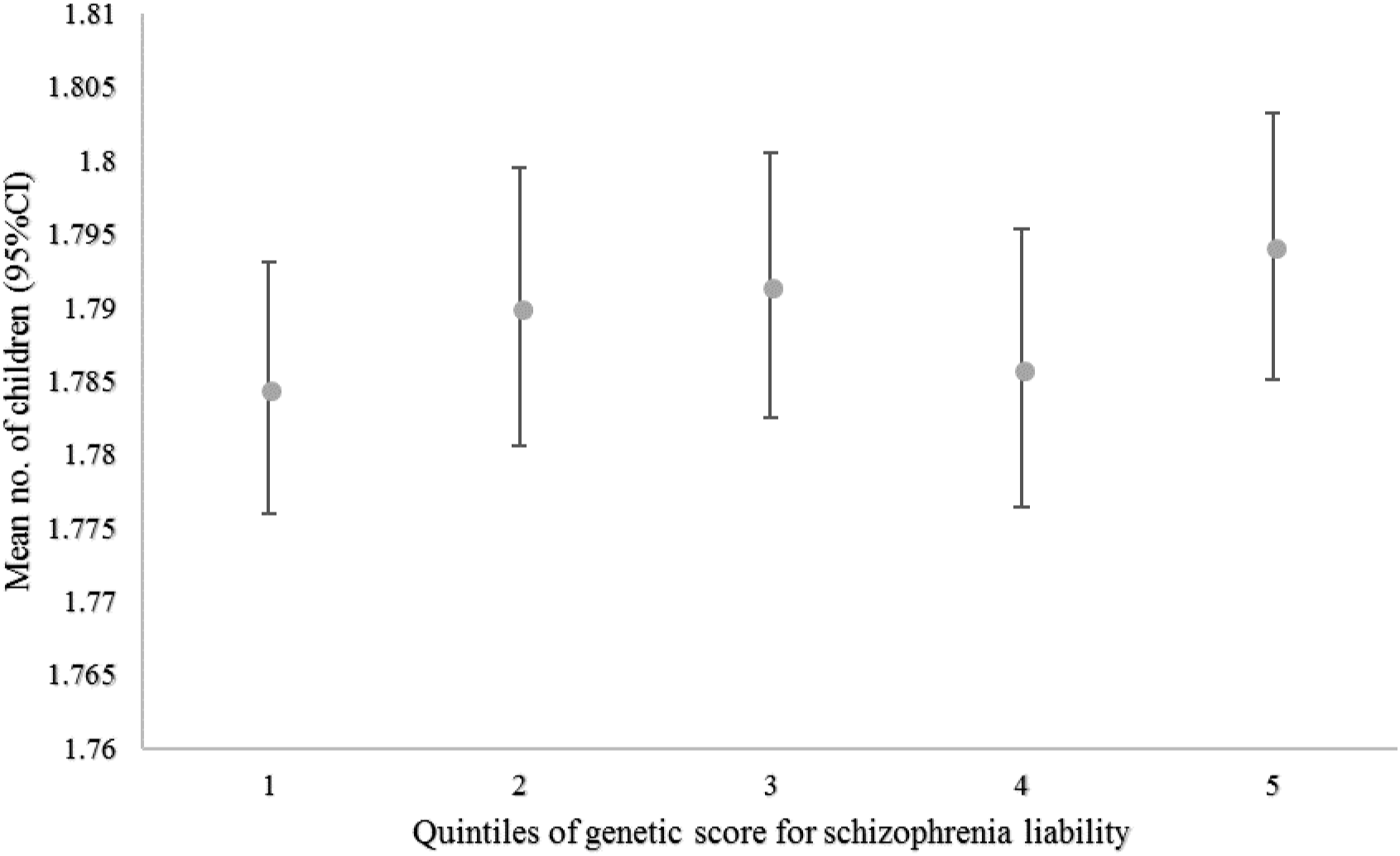
Genetic score for schizophrenia liability (in quintiles) and mean number of children in UK Biobank data showing little evidence of heterogeneity across values of the schizophrenia score.

**Figure 2.**
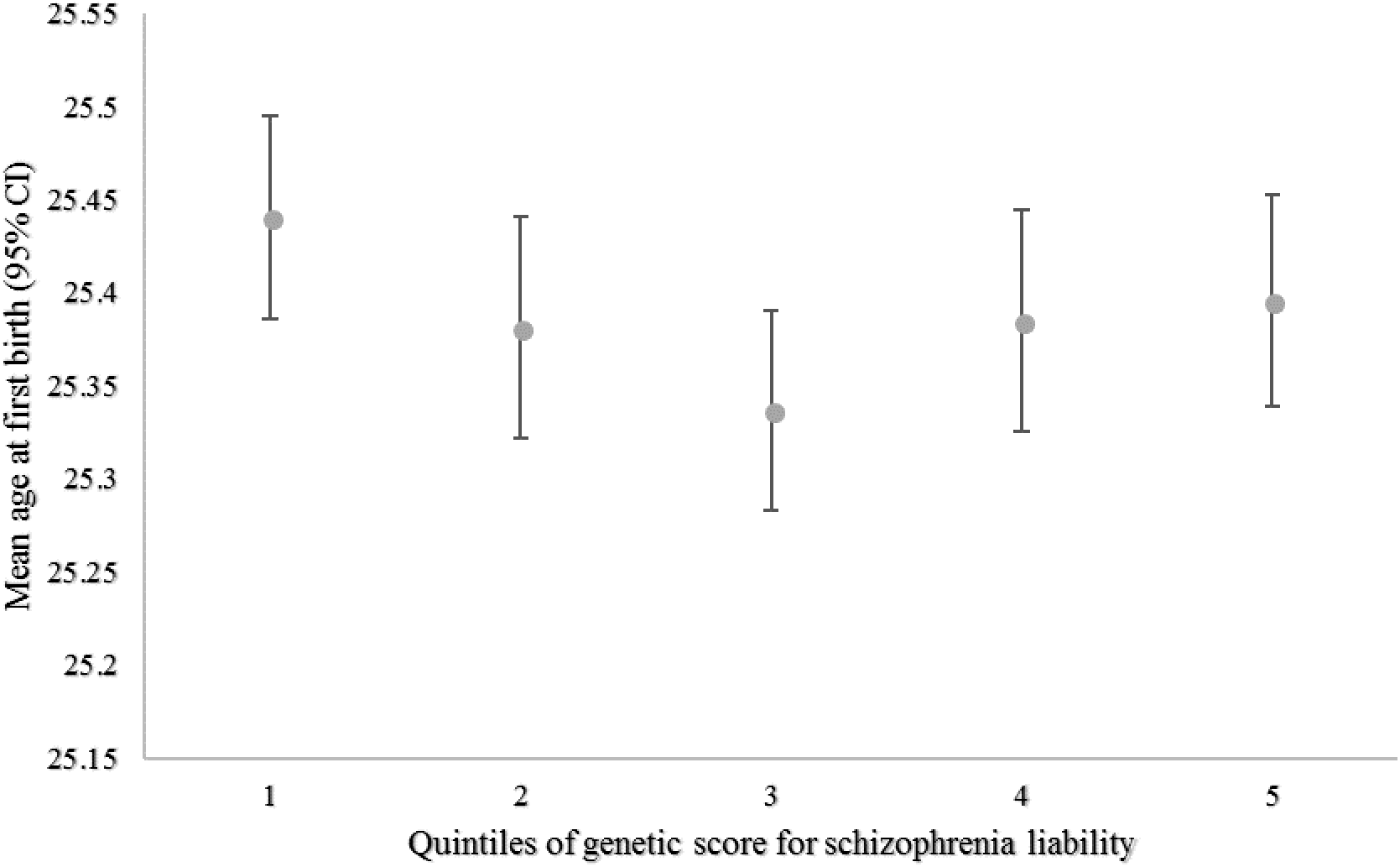
Genetic score for schizophrenia liability (in quintiles) and mean age at first birth in women from UK Biobank data also showing little evidence of heterogeneity across values of the schizophrenia score.

**Table 3.** Associations of the score for genetic liability for schizophrenia and number of children removing cumulative deciles of the score. Adjusted for the top 10 principal components.

| Schizophrenia genetic score | Number of children |  |
| --- | --- | --- |
| | *N* | $\beta$ (95% CI), *P* |
| Highest 10% removed | 302,190 | 0.0005 (-0.0003, 0.0013), 0.19 |
| Highest 20% removed | 268,604 | 0.0002 (-0.0007, 0.0011), 0.70 |
| Highest 30% removed | 235,030 | 0.0005 (-0.0006, 0.0016), 0.35 |
| Highest 40% removed | 208,433 | 0.0006 (-0.0006, 0.0019), 0.30 |
| Highest 50% removed | 167,860 | 0.0008 (-0.0006, 0.0023), 0.27 |

## Discussion

Our results do not indicate a genetic correlation between genetic liability for schizophrenia and reproductive success using LD score regression, or a linear causal effect between these using MR techniques. This is inconsistent with cliff edge-fitness maintaining schizophrenia in the population, which would predict an increase in fitness with increased genetic liability in the general population. These results support previous research suggesting no strong evidence of a relationship between genetic liability for schizophrenia and number of offspring (8,25). Also consistent with previous research, we found no clear evidence of linear association between genetic liability of schizophrenia and age at first birth (8). In sensitivity analyses, we found some suggestion of a possible peak in fitness at intermediate to high levels of genetic liability, but there was no statistical evidence for this, suggesting that if this non-linear association exists it is very weak, and not reliably detectable even in a large study such as UK Biobank. A previous study also showed little evidence of quadratic associations between genetic liability for schizophrenia and reproductive success (8). Results of our positive control analyses were as expected and in line with previous genetic research, suggesting that educational attainment is under negative selection (25–29), which suggests that the overall approach we adopted here is valid.

Cliff-edge fitness suggests that schizophrenia prevalence is sustained because the negative reproductive effects in those with an underlying genetic liability and the disorder are offset by a reproductive advantage to those who have an underlying genetic liability but do not develop the disorder (8). We therefore only examined part of the cliff-edge hypothesis by studying only those without the disorder, testing whether there is a linear effect on fitness with increasing genetic liability. Although it is hard to estimate the size of effect on fecundity necessary to sustain the prevalence of schizophrenia (or indeed whether this effect size may fall within the confidence intervals of our estimate), our data provide no evidence in support of a cliff edge fitness effect. This leaves us with two alternative theories for how schizophrenia prevalence is maintained. One is that as schizophrenia is a highly heterogenous disorder and exhibits a highly polygenic architecture, and effects of genetic variants are individually too weak to be under negative selection (1,8,10). Our results are consistent with this possibility and suggest that identified schizophrenia risk variants are not under strong selection in the general population. Another explanation is that mutation-selection balance maintains the prevalence of schizophrenia; rare recurrent DNA copy number variants which are also risk factors for schizophrenia are filtered out of the population by selection and replenished by *de novo* mutations (9). Rare copy number variants conferring risk to psychiatric illness are under strong negative selection (8,9), with most persisting in the population for only two generations (9). We used results from GWAS, which mainly detect common alleles and therefore cannot determine whether mutation-selection balance sustains the prevalence of schizophrenia through rare variants, although rare variants have been shown to associate with number of children (1,8). Other explanations could include an increased likelihood of symptom diagnosis, changes in the enviornment (52,53) and/or selection bias. UK Biobank data is unrepresentative of the population, given a response rate of approximately 5%, which may introduce selection bias (32,54). This can generate spurious results in genotypic associations when selection is based on phenotypes associated with the genetic variants and could attenuate associations towards the null if schizophrenia-proneness and increased number of children reduced participation (55–57). Previous studies have found that higher genetic liability for schizophrenia is associated with lower participation in cohort studies which could bias estimates between genetic liability and traits that lead to nonparticipation in genetic associations and MR (56,58).

A key strength of this study is the use of MR, which can provide stronger evidence of causality than observational studies (18,59). We showed agreement between various MR methods that rely on differing assumptions and agreement between methods, which provides greater confidence in the robustness of the results (60). We further conducted a positive control analysis to confirm that our approach was valid. Additionally, our MR offers large sample sizes which are necessary for investigating small effect sizes common in such genetic analysis (43). However, there are also some limitations that should be considered with the current evidence. Firstly, MR relies on genetic variants naturally randomizing an exposure, and therefore inferring causality from genetic liability for schizophrenia as the exposure requires careful interpretation. Our outcome sample was not selected on schizophrenia status, so it contained only few cases of diagnosed schizophrenia. Therefore we assume that schizophrenia SNPs are associated with sub-diagnostic schizophrenia traits that could cause a reproductive advantage within the wider population (4,13,15). Although debated (61,62), schizophrenia symptoms have been suggested to exist on a continuum, and this assumption could therefore be met (61,63–65). Within this, we assume that the instrumental variable assumptions are satisfied for this continuous liability to provide a valid test of causality using the binary exposure (49). Secondly, variants are non-specific and it is difficult to fully remove population structure which can induce spurious associations through confounding, even within a sample of European ancestry and adjusting for principal components as we have done (30,66). Lastly, age at first birth was only measured in females in UK Biobank, and therefore consists of a different population to the exposure data (which includes data from both females and males). However, the correlation between male and female estimates for age at first birth in a recent GWAS was high (28).

The present study highlights the continued importance of investigating differential fertility and contributes to understanding the maintenance of schizophrenia, and educational attainment, in the population (3,22,67). Educational attainment has previously been shown to predict human longevity (68) and highlights how even traits with a positive effect on longevity can be maladaptive, although other influences on educational attainment in the population are also identified (29). This work additionally demonstrates how epidemiological methods can be repurposed to study evolutionary theories. Future research should investigate causal methods for estimating non-linear relationships as well as other explanations for this evolutionary paradox, such as mutation-selection balance.

## Acknowledgments

RBL, HMS, AET, REW, GDS, NMD, GH, AF and MRM are members of the Medical Research Council Integrative Epidemiology Unit at the University of Bristol, which is supported by the Medical Research Council and the University of Bristol and funds RBL’s PhD studentship (grant number: MC_UU_00011/7). AF is supported by a personal fellowship from the UK Medical Research Council (MR/M009351/1). AET and MRM are members of the UK Centre for Tobacco and Alcohol Studies, a UKCRC Public Health Research: Centre of Excellence. Funding from British Heart Foundation, Cancer Research UK, Economic and Social Research Council, Medical Research Council, and the National Institute for Health Research, under the auspices of the UK Clinical Research Collaboration, is gratefully acknowledged. GH is funded by the Wellcome Trust (208806/Z/17/Z). The Economics and Social Research Council (ESRC) support NMD via a Future Research Leaders grant [ES/N000757/1]. REW, ISP-V and MRM are supported by the NIHR Biomedical Research Centre at the University Hospitals Bristol NHS Foundation Trust and the University of Bristol. This research has been conducted using the UK Biobank Resource under Application number 6326. The authors also thank Dr Suzi Gage, Dr Jack Bowden and Dr Dan Lawson for their technical support and comments. The authors also thank Dr Ruth.Mitchell, Dr Gibran Hemani, Mr Tom. Dudding and Dr Lavinia. Paternoster for conducting the quality control filtering of UK Biobank data. The authors are grateful to the participants of UK Biobank and those who contributed to the PGC and SSGAC GWAS, as well as research staff who worked on the data collection.

## Ethical statement

UK Biobank received ethics approval from the Research Ethics Committee (REC reference for UK Biobank is 11/NW/0382).

## Data accessibility

Genome wide summary data for schizophrenia and educational attainment can be downloaded from the Psychiatric Genomics Consortium and Social Science Genetic Association Consortium websites (http://pgc.unc.edu; https://www.thessgac.org/). UK Biobank data is available upon application (www.ukbiobank.ac.uk). Analysis scripts are available on GitHub (https://github.com/MRCIEU/Schizophrenia_Fertility_Paper.git).

## Author contributions

Contributors AF, MRM, and ISP-V conceived the study. RBL conducted the analysis and drafted the initial manuscript. HMS, REW, GH, GDS, NMD and AET assisted with analysis and interpretation. All authors assisted with interpretation, commented on drafts of the manuscript and approved the final version.

